# Bridging the gap with invasive imaging: promises and challenges of a new generation of ultrahigh resolution fMRI

**DOI:** 10.64898/2026.04.10.717824

**Authors:** Lasse Knudsen, Yulia Lazarova, Steen Moeller, Nils Nothnagel, Lonike K. Faes, Essa Yacoub, Kamil Ugurbil, Luca Vizioli

## Abstract

The human neocortex is organized into six laminae forming the structural basis for feedforward and feedback connections across the brain, yet their functional contributions have remained largely inaccessible for non-invasive imaging methods. Leveraging the ultrahigh field of a 10.5 Tesla scanner, we acquired anatomical and functional MRI data at 0.37mm (∼0.05 µL) and 0.35mm (∼0.04 µL) isotropic resolution, respectively, approaching the scale of individual cortical layers in humans. Using the Stria of Gennari as an in-vivo anatomical landmark, we extend our previous finding that feedforward visual activation in layer IV of the primary visual cortex during visual stimulation was resolved in laminar BOLD profiles. These laminar features were reproducible across sessions and were not clearly visible with more typical 0.8 mm resolutions at 7T, underscoring the benefits of further increases in magnetic field strength and resolution. This imaging domain, however, comes with increasing challenges of distortion, alignment, and cortical depth estimation, which must be addressed and mitigated to realize its benefits. In this paper we discuss the promises and challenges of this new regime of high resolutions. Our findings showcase the potential of ultrahigh field, ultrahigh resolution human fMRI to bridge the gap with invasive imaging of cortical layers.

## Introduction

Since its introduction in 1992^1,2^, blood oxygen level-dependent (BOLD) functional magnetic resonance imaging (fMRI) has become one of – if not the – most powerful tool to study the human brain non-invasively. Recent technological advances in ultrahigh-field MRI, paired with the growing availability of ultrahigh field systems (≥7T), have increased the spatial precision of such studies from millimeter to submillimeter spatial resolutions, initiating efforts to penetrate the mesoscopic scale such as cortical layers^3–5^.

As established by research using animal models, cortical layers are characterized by different cell types and concentrations with layer-specific connectivity patterns: in primary sensory areas, feedforward signals predominantly target middle layers, whereas feedback connections terminate in inner and outer layers^6,7^. Brain function relies on this layer-dependent connectivity, combining them into recursive circuital interactions, crucial for key human processes such as perception, memory and consciousness. Resolving functional mapping at the mesoscopic scale is thus essential to understand the human brain not only in healthy individuals but also in patient populations. Perturbations in these processes have been proposed to model dysfunction in widespread neuropsychiatric and neurodegenerative diseases, including schizophrenia, Parkinson’s, and Alzheimer’s disease^8–13^.

However, reliably measuring layer-dependent computations with fMRI remains challenging. To date, most human laminar fMRI studies have employed ∼0.8 mm isotropic voxels (∼0.5 µL), a resolution that has revealed key principles of the human brain’s laminar function across diverse domains, including visual^14–18^ and auditory perception^19,20^, somatomotor function^13,21–25^, language^26^, axis of motion^27,28^, and working memory^29,30^. Yet, given that cortical thickness ranges between ∼1 and 4.5 mm^31^, voxel sizes on the order of ∼0.8 mm may not suffice to fully capture subtle but functionally meaningful modulations across layers^32,33^, depending on the brain region of interest and the paradigm used. This limitation is further compounded by a multitude of factors known to degrade the functional precision of echo planar imaging (EPI) readouts, resulting in an effective resolution below the nominal value^34,35^. To more accurately characterize fine-grained features of laminar activation in the human brain, approaching those achieved in animal fMRI studies^36–40^, substantially higher resolution and functional contrast-to-noise (fCNR) are required (as illustrated in^41^). This was already anticipated in the strategic plan for the BRAIN initiative,^42,43^ which challenged the field to reach spatial resolution with <0.01 µL^44^ voxel volumes in humans, approaching the scale of individual cortical layers. Such resolutions need to be attainable at the single-subject level, to enable individualized computational modelling^45–48^ and patient-specific evaluation of neuroprogressive diseases known to be layer-specific^8–13^.

Leveraging the supra-linear gains in signal-to-noise ratio (SNR), the BOLD effect, and fCNR with higher field strengths^49–52^, combined with NORDIC denoising^53,54^ and in-house built high channel count RF arrays^49,51,55^, we recently explored the feasibility of robust functional acquisitions at 0.35 mm isotropic (0.042 µL) resolution in well-trained human subjects at 10.5 T^56^. With this resolution, representing a 12-fold volume decrease relative to the typically employed ∼0.8 mm isotropic voxels^56^, each gray matter voxel contains on the order of <2000 neurons^42,57^, approaching the neuronal sampling capability of invasive optical imaging. To enhance fCNR, which is limiting at these resolutions, we employed a gradient-echo BOLD (GE-BOLD) EPI acquisition protocol. GE-BOLD is traditionally criticized for its large draining vein weighting, compromising spatial specificity^58–61^. However, this limitation is alleviated precisely in the ultrahigh field and ultrahigh resolution regime. At ultra-high fields, we are increasingly sensitive to the microvasculature, and at ultra-high spatial resolution, the T2*-dephasing around large vessels–the root cause for the loss in spatial specificity–decreases, with a consequent reduction of the contribution coming from large vessels (see^53,56,62^ and *Discussion*). These factors allow the superior sensitivity of GE-BOLD to be exploited with reduced impact of large veins, making it well suited for the pursuit of non-invasively resolving individual cortical layers.

In this work, we discuss the needs, promises and challenges of anatomical and functional acquisitions at resolutions capable of resolving individual cortical layers, thereby narrowing the gap between human fMRI and invasive animal studies. Specifically, using GE-BOLD at 0.35 mm isotropic resolution and 10.5 T, we demonstrate the reliability of our previous finding that a feedforward-driven activation peak aligned with layer IV (Brodmann classification) in V1 could be captured in laminar BOLD profiles during visual stimulation^56^. At this resolution, layer IV could be identified directly in functional space from time-series-averaged EPI intensities using the Stria of Gennari (henceforth referred to as Stria) as a landmark^63–66^, validated against an independent anatomical reference scan. We discuss our initial experiences with this new generation of ultrahigh field/ultrahigh spatial resolution fMRI data, highlight arising challenges of distortion, alignment, and cortical depth estimation and provide potential solutions.

## Results

We acquired functional images at 10.5 T using a 3D GE-EPI sequence during a block design visual stimulation paradigm with flashing target/surround checkerboards (see *Methods*). Statistical analyses were performed at the single-subject level, each representing a test-retest case, with statistical power derived from the number of single trials. We additionally acquired a whole-brain 0.37 mm isotropic resolution multi-echo GRE (ME-GRE) scan, used to support segmentation, and for identifying the Stria (see *Methods*).

### Imaging cortical layers at ∼0.04 µL resolution

Figure 1A shows example slices from the ME-GRE anatomical reference scan and from the mean-EPI data (averaged across individual volumes within a single fMRI run). The Stria appears as a line of reduced intensity along the cortical ribbon of V1 in the ME-GRE image (blue arrows in the zoomed-in insets), consistent with previous results^65^, and is also discernible on the EPI image. In the laminar profiles, this landmark manifests as a dip in the T2*-weighted intensity values (blue arrow in the profile plot, Figure 1A), thereby identifying layer IV. Functional time series were NORDIC denoised for thermal noise suppression, making the Stria visible even in single functional images (Figure 1B; see also Supplementary Fig. 1).

**Figure 1.**
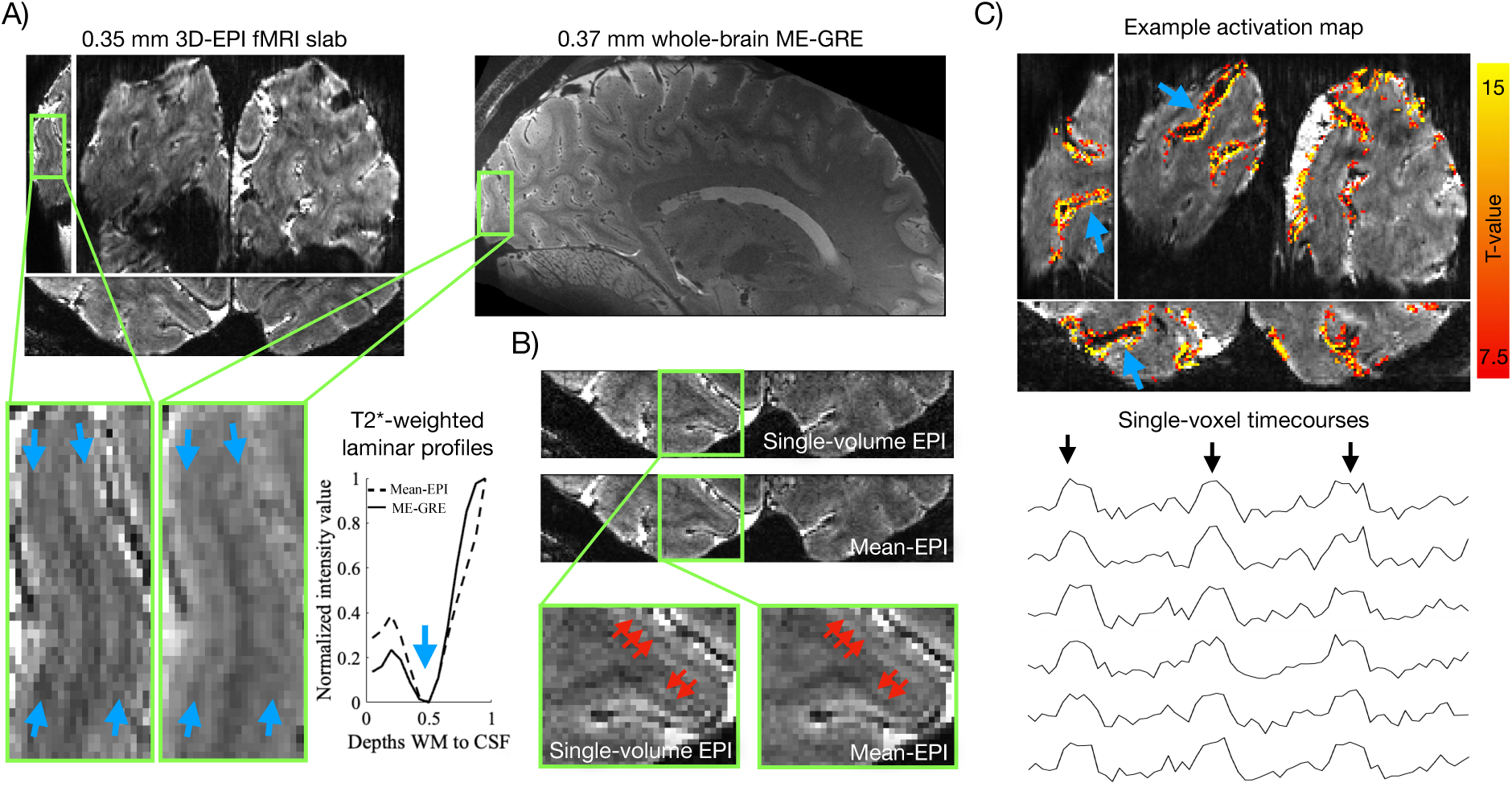
Anatomical and functional acquisitions at ∼0.04µL resolution. **A)** Mean-EPI and ME-GRE images from an example subject. Blue arrows in the zoomed insets highlight the Stria of Gennari, visible in both anatomical and functional data. Corresponding laminar profiles reveal a dip in normalized intensity values at the depth of the Stria (blue arrow). **B)** Example single-volume EPI and mean-EPI images. The offline reconstruction (see *Tailored offline reconstructions* in *Methods*) yielded high SNR and reduced blurring (see Supplementary Fig. 1 for comparison with default vendor reconstructions), allowing visualization of the Stria even in single-volume EPI images (highlighted by red arrows). **C)** Upper panel: example activation maps (t-values, target > fixation) with blue arrows indicating separation of activation across adjacent cortical banks. Lower panel: example high-CNR single-voxel, single-run time courses showing signal increases aligned with target stimulation blocks (black arrows).

Figure 1C (upper panel) displays single-condition t-value activation maps from an exemplary subject during flashing checkerboard stimulation (target > fixation; maps from the remaining subjects are shown in Supplementary Fig. 2). Robust activation was observed with ∼15 minutes of data, indicating strong BOLD sensitivity. This is further supported by single-voxel, single-run time courses exhibiting clear magnitude changes co-occurring with the stimulation blocks (Figure 1C, lower panel; Supplementary Fig. 2). The activation followed the cortical ribbon and was clearly separable between adjacent banks of corresponding sulci (exemplified by blue arrows), consistent with high spatial specificity.

Figure 2A (upper panel) shows the BOLD single-condition laminar profiles (percent signal change (PSC)) elicited by the target stimuli in a region of interest (ROI) within the target-representation of V1 (example slice shown in the lower panel of Figure 2A; details outlined in *Methods*). In line with expectations of a pure feedforward stimulation paradigm as used here^36–38,63,67,68^, the profiles of all subjects exhibited a significant local activation peak (p<0.05, see *Methods* and Supplementary Fig. 3) co-localizing with the Stria dip (i.e. layer IV) in the corresponding mean-EPI profile (Figure 2A, middle panel), thereby replicating and expanding upon our recent findings^56^ in four new datasets. Activation at the Stria was also evident directly in activation maps and was reproducibly observed across sessions in two subjects scanned twice (Subject 1 and Subject 2; Figure 2B, smoothed within layers to enhance laminar features using LAYNII^69^, *LN2_LAYER_SMOOTH*-function, FWHM=2 mm).

**Figure 2.**
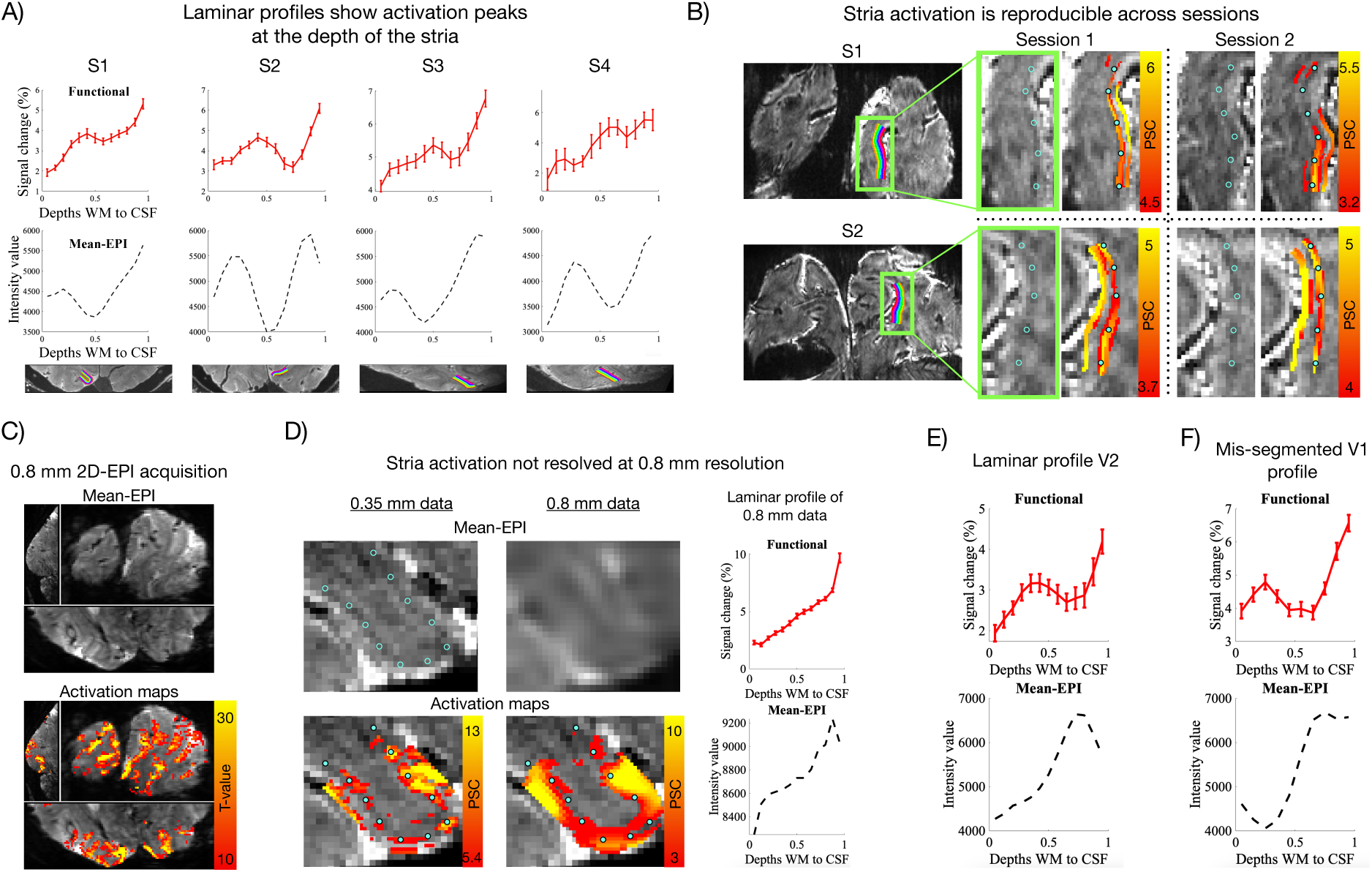
Resolving functional activation peaks at the Stria. **A)** Functional (upper panel) and mean-EPI laminar profiles (middle panel) from all subjects (S1-S4), extracted from the target representation of V1 (example slice shown below), showing functional peaks aligned with the dip in the mean-EPI profile marking the Stria. **B)** Reproducibility across sessions in two subjects (S1, S2), with activation (PSC) consistently falling on the Stria; maps were smoothed within layers and thresholded to enhance visibility of laminar features. Blue rings indicate the Stria on the mean-EPI underlay and are copied as circles onto activation maps as a reference. **C)** Mean-EPI and activation maps (t-values, target > fixation) from a 7T/0.8 mm acquisition. **D)** Comparison of 0.35 mm versus 0.8 mm data, showing that at 0.35 mm the Stria traversed the middle of the GM ribbon with activation falling on top, whereas at 0.8 mm neither the Stria nor a corresponding activation peak was resolved, as confirmed by the adjacent laminar profiles. **E)** Laminar profiles from V2, showing a functional peak without a corresponding decrease in intensity values in the mean-EPI profile in V2. **F)** Laminar profiles from V1 with a deliberately misplaced WM/GM boundary (omitting deepest depths), illustrating how genuine layer IV activation, validated by Stria overlap, could be misattributed to deep layers without the Stria landmark. Error bars in functional profiles represent standard error of the mean across trials.

To provide a reference for a more conventional laminar fMRI setup, Subject 1 was additionally scanned at 7T using 0.8 mm isotropic voxels with the same visual stimulation paradigm (Figure 2C-D, see^56^ for more such datasets). At this lower resolution and field strength, the Stria was no longer clearly identifiable neither in the mean-EPI image (Figure 2C and zoomed inset in Figure 2D) nor in the corresponding laminar profile (Figure 2D). Similarly, the local BOLD activation peak was not resolved (Figure 2D; p>0.05, see Supplementary Fig. 3), most likely due to increased partial voluming and/or higher sensitivity to large veins (see *Discussion*). Such vascular contributions are known to displace functional BOLD signals from the underlying neuronal source, thus inducing a superficial bias^58–61^. This effect is evident as a monotonic increase in BOLD signal toward the cortical surface (maps in Figure 2C-D, profile in Figure 2D), and has been consistently reported in the literature^56,70–75^.

Figure 2E shows the laminar profile obtained from V2 in an example subject (Subject 1). A significant activation peak was also observed in the middle of the cortex in V2 (p<0.05, see Supplementary Fig. 3). Importantly, mean-EPI in this region does not exhibit decreases in intensity values in the middle of the cortical ribbon comparable to that observable in V1 corresponding to the highly myelinated Stria. This result is compatible with the notion that the observed V1 functional peaks cannot be explained solely by scaling related to mean-epi intensity values (see discussion).

The absence of laminar landmarks (such as the Stria), as is the case at 0.8mm isotropic resolution (Figure 2D), holds the risk of ambiguous interpretation as illustrated in Figure 2F. Here, the white matter (WM)/gray matter (GM) boundary in V1 was deliberately superficially misplaced. Activation appeared to peak in the deep layers, which would suggest feedback-driven input. However, with the Stria-dip in the mean-EPI profile serving as a guide, this peak can be (correctly) assigned to layer IV activation, suggesting feedforward-driven input.

### Resolving laminar features requires consistent layer positioning

The accuracy of the alignment of functional peaks in laminar profiles to the underlying biological layers, is determined by layer positioning— the correspondence between estimated cortical depths/layers and biological cytoarchitectonic layers—both within and across slices. Layer positioning is determined by segmentation boundaries, layer-estimation approach (equidistance versus equivolume^76^), and algorithmic parameters such as the number of smoothing iterations^69^ (Figure 3A).

**Figure 3.**
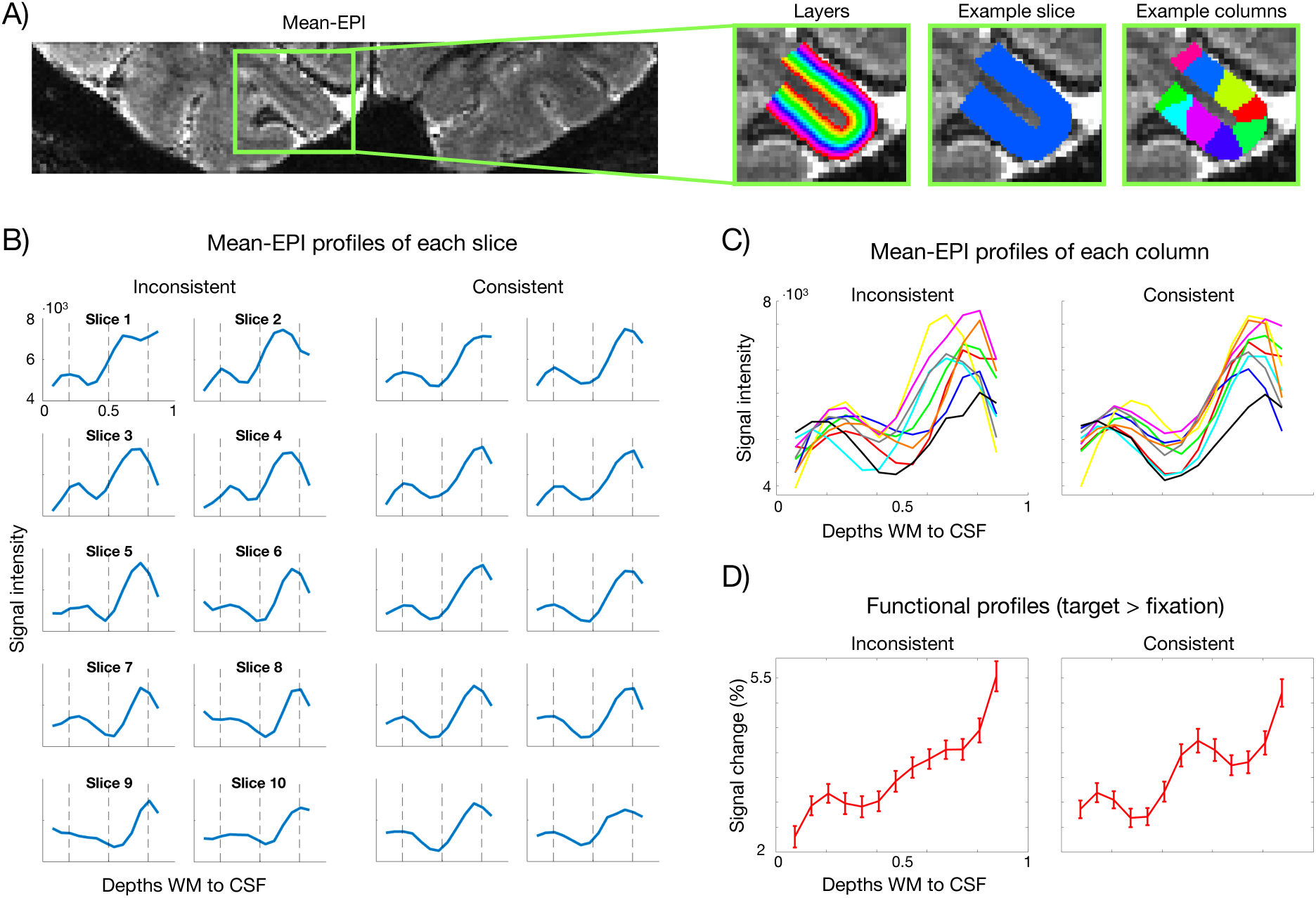
Consistent layer positioning is critical at ultrahigh spatial resolution. **A)** Depiction of layers, as well as example slice and column masks used for plotting laminar mean-EPI intensity profiles to ensure consistent layer positioning. **B-C)** Laminar EPI intensity profiles from each slice and column in an example ROI, before (inconsistent) and after (consistent) correction of layer positioning based on these plots. **D)** Functional laminar profiles averaged across the ROI. The functional peak at the stria-dip emerged with consistent layer positioning. Error bars in functional profiles represent standard error of the mean across trials.

At high resolution in particular, inconsistent layer positioning may result in laminar features being mixed/blurred when averaging across the ROI, thereby reducing laminar specificity. To this end, the visibility of anatomical layer-specific landmarks, such as the Stria, allows us to assess and correct the layer positioning.

ROIs were subdivided into slices and columns (Figure 3A) and mean-EPI profiles were plotted for each (see *Methods* for details). To illustrate the impact of alignment, we show an example where laminar features in the profiles, hereunder the Stria dip, were misaligned (inconsistent, before correction) versus aligned (consistent, after correction) across slices and columns (Figure 3B–C). Notably, despite the average shift in the Stria being only ±0.25 mm (SD=0.08 mm) across slices for this exemplar inconsistent layer positioning (see *Methods*), the resulting functional profiles averaged across the ROI (Figure 3D) clearly revealed the expected layer IV peak only under consistent alignment, exemplifying the critical role of precise layer positioning at ultrahigh resolution. To further support this notion, we carried out a supplementary analysis in which depth maps were systematically shifted in opposite directions (with respect to activation maps) across slices to mimic intra-ROI layer inconsistencies (see *Methods*). The smallest shift size resulting in a significant difference between shifted and unshifted profiles was ±0.24 mm (0.48 mm total shift between slices) on average across subjects (Supplementary Fig. 4).

### Alignment challenges in the regime of ultrahigh field and resolution

Geometric distortions were pronounced in the data presented here, as a consequence of the higher field strength and long readout times associated with high-resolution acquisitions^77,78^. Figure 4A illustrates this by showing two EPIs acquired with the same sequence but opposite phase-encoding directions (the blue contour indicates the outline of the forward scan). Such distortions complicate co-registration with minimally distorted anatomical reference scans, such as the ME-GRE scan used here. Phase-reversed distortion correction (Blip-up/blip-down)^79,80^ corrected distortions at the global level, yet, local discrepancies remained (highlighted by the red and green contours tracing local structures of the forward and reverse scans, respectively; Figure 4B). Co-registration of the structural scan to the EPI (see *Methods*) tended to fail in these same regions of severe residual distortion, even with non-linear transforms (Figure 4C, red and green contours outline EPI and anatomical structures, respectively). In contrast, accurate co-registration—to the precision of the Stria—was feasible in regions where residual distortions were small, and image contrast was high (zoomed insets in Figure 4D; an illustration of the minimal residual distortions in the zoomed region is available in Supplementary Fig. 5).

**Figure 4.**
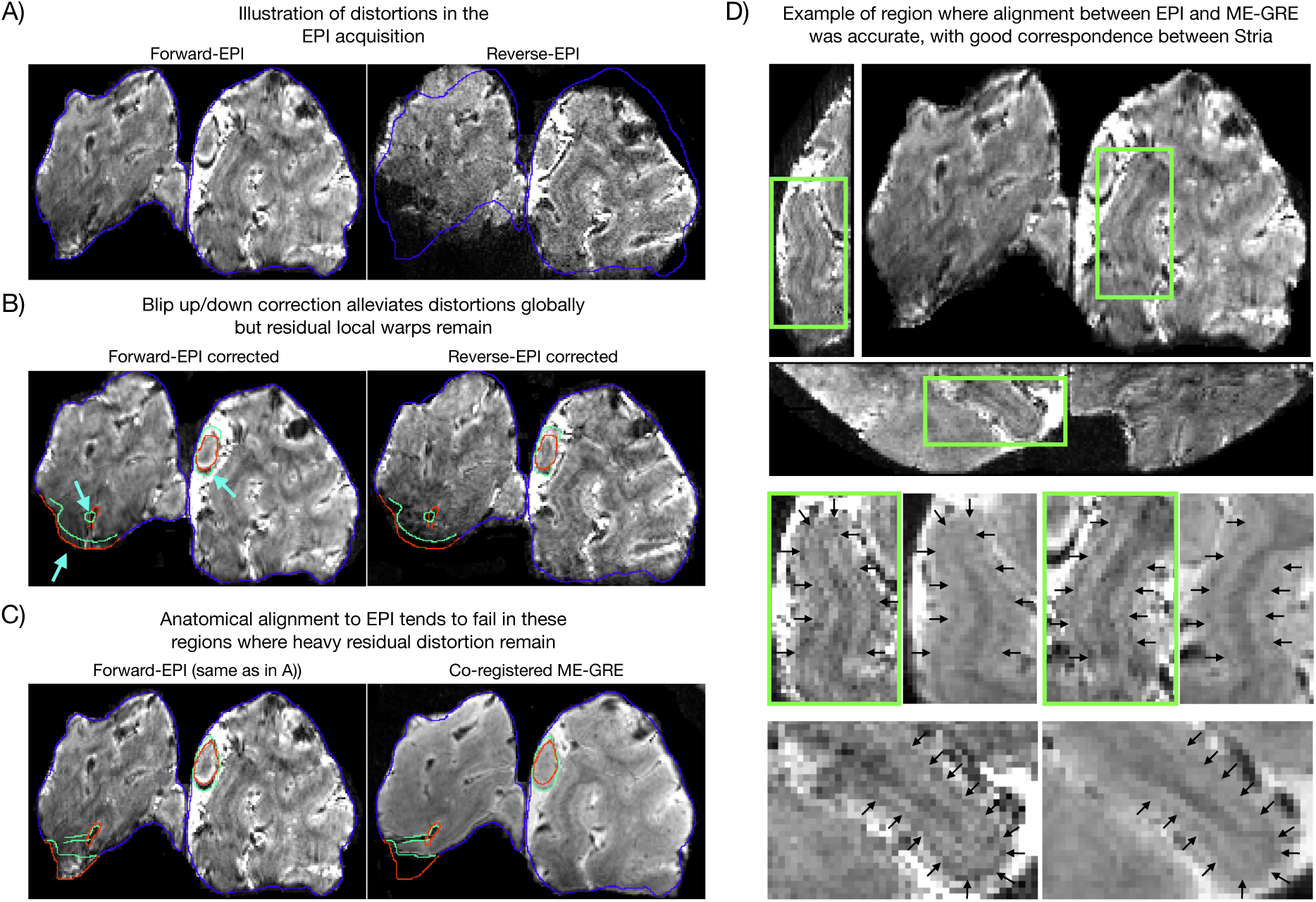
Co-registration challenges at ultrahigh field and resolution. **A)** Example coronal slice showing forward-EPI (foot-head) and reverse-EPI (head-foot) acquisitions. Spatial distortions are pronounced, as seen in the mismatch of the reverse-EPI within the blue outline drawn on the forward-EPI. **B)** Global mismatch was reduced after distortion correction, but significant residual distortions remained locally, highlighted by red and green contours tracing structures in the forward and reverse scans, respectively. **C)** Co-registration between ME-GRE and mean-EPI images tended to fail in regions where severe residual distortions remained (see red and green contours outlining EPI and anatomical structures, respectively). **D)** Example of a region with accurate registration, minimal residual distortions and high image contrast. Black arrows indicate Stria overlap across modalities (i.e. the mean-EPI and the ME-GRE).

Motion correction in most fMRI software packages is typically performed using 6-parameter linear estimation (3 translations and 3 rotations), which assumes rigid-body displacements only. However, distortions are dynamic across an imaging session^77,81^. Given the pronounced distortions at 10.5 T and the increased need for alignment precision at 0.35 mm resolution, we evaluated whether non-linear alignment would be beneficial.

Figure 5A compares the accuracy of linear versus non-linear registration across runs, where the mean of each run was registered to the mean of a reference run (see Methods). Accuracy was quantified by the overlap of z-scored single-run mean-EPI profiles, expressed as the root mean square (RMS) error relative to the overall mean-EPI profile. The RMS error was significantly lower with non-linear across-run alignment in three of four subjects (two-sided paired t-tests, Figure 5A); in the remaining subject, no significant difference was observed. For within-run motion correction, however, we refrained from non-linear alignment as it has been suggested to potentially confound activation-induced signal changes with motion and thereby suppress BOLD signal (talk at ISMRM 2024, “Ultrahigh spatial resolution imaging in the presence of motion”, Laurentius Huber). This was also observed in our data as shown in example activation maps and time series (Figure 5B). Overall, linear within-run motion correction produced consistent alignment between volumes based on both overlap of single-volume profiles (Figure 5C) and visual inspection of motion videos (Supplementary Fig. 6). In Subject 4, two runs were exceptions to this with large RMS-errors of single-volume EPI profiles (green box in Figure 5C). These runs were notably characterized by extreme motion relative to the voxel size (>2 mm within-run translations). The functional laminar profile of these runs is shown in Figure 5D (green), which deviated markedly from the expected pattern (i.e. peak at the Stria) observed in the remaining runs (black; Figure 5D).

**Figure 5.**
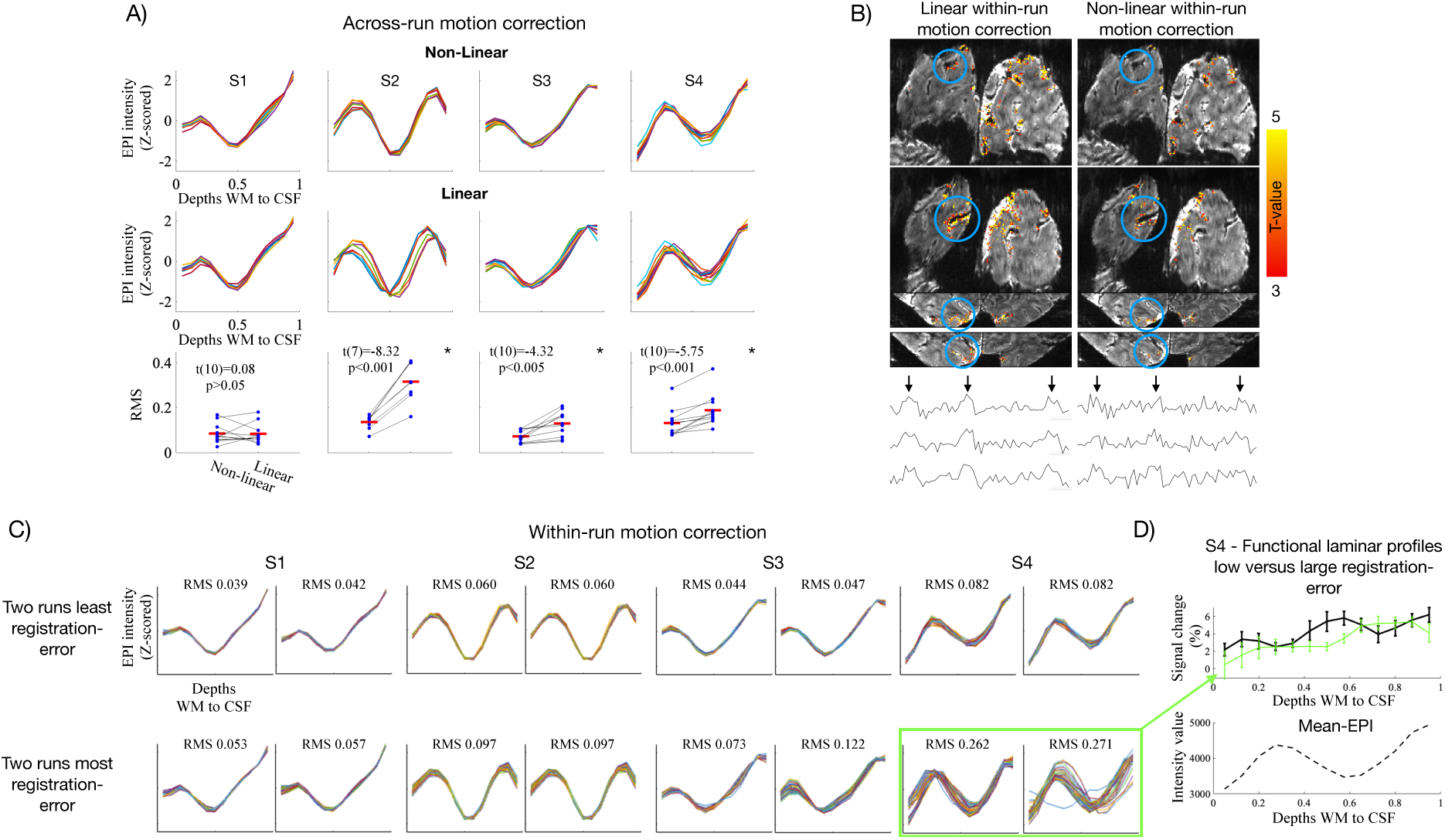
Evaluating non-linear motion correction across and within runs. **A)** Single-run mean-EPI profiles (z-scored) for each subject, aligned using linear + non-linear (upper panel) or linear-only (middle panel) transforms. Lower panel shows RMS error of each run relative to the across-run mean. Mean RMS (red bars) was significantly higher for linear-only transforms in three subjects and unchanged in the fourth. **B)** Activation maps (t-values) from a single run motion-corrected with linear-only or linear + non-linear transforms. Blue circles mark regions of signal suppression after non-linear correction, as also seen in example single-voxel time courses (black arrows indicate target stimulation blocks). **C)** Single-volume EPI laminar profiles illustrating within-run alignment under linear correction, shown for the two lowest and two highest RMS-error runs per subject. Green box highlights the two runs with the largest RMS error across subjects. **D)** Functional laminar profiles for the two high-RMS-error runs (green) and the remaining runs (black). Error bars in functional profiles represent standard error of the mean across trials.

## Discussion

In the present study, we demonstrate the feasibility of ultrahigh field and resolution GE-BOLD fMRI data to co-localize with layer IV in human visual cortex. In the following sections we discuss these findings and address the challenges inherent to this regime and potential strategies to overcome them.

### The potential of higher spatial resolution fMRI

As outlined in the *Introduction*, non-invasive imaging methods capable of accessing cortical layers are essential for understanding mesoscale computations constituting the cortical foundation of human brain function^7,82,83^ and neurodegenerative disorders^8–10^. To this end, the advent of laminar fMRI has sparked excitement and has already proven its value in both basic neuroscience^14–20,22–26,29,30,84^ and clinical research^8,11–13,85^. However, most laminar fMRI studies, typically employ ∼0.8 mm isotropic voxels, which may be too coarse for resolving intricate laminar responses.

The present study demonstrates the potential of achieving higher spatial resolution fMRI, exploiting the increased SNR and BOLD effect of very high magnetic fields, in this case 10.5T. We extend our recent finding^56^ that GE-BOLD at 0.35 mm isotropic resolution could resolve the expected feedforward-driven activation peak in layer IV of V1^36–38^ at the single-subject level (Figure 2A-B, Supplementary Fig. 3). It should be noted that since PSC is computed as a ratio of response amplitudes relative to its mean EPI baseline, increases in PSC amplitudes may be related to decreases in local mean EPI intensity values associated with the Stria dip in V1 (Figure 2A). Here, however, we report that a clear local activation peak away from the surface was also detectable in V2, despite the absence of the local decrease in baseline EPI images (Figure 2E, Supplementary Fig. 3). This result is compatible with the notion that the reported activation peak is of neuronal origin.

The layer IV peak is generally not detectable with GE-BOLD EPI at ≤7T and ∼0.8 mm isotropic voxels (Figure 2C-D, Supplementary Fig. 3, see also^56,70–75^), due to partial volume effects combined with the limited spatial specificity of GE-BOLD related to its sensitivity to draining veins (discussed in *Sensitivity to large draining veins*)^58–61^. FMRI activity peaking away from the surface towards the middle depths of V1 has been observed previously with non-GE-BOLD acquisitions that possess higher microvascular specificity, such as VASO, spin-echo BOLD or ASL^70–73,75,86,87^. However, these come with significant reduction in fCNR and temporal efficiency, constraining their sensitivity, coverage and spatiotemporal resolution. As such, GE BOLD remains the method of choice in functional brain mapping in humans. The methodological advance of being able to resolve such localized responses with ultrahigh resolution GE-BOLD, is three-fold: 1) this acquisition strategy offers high readout efficiency, supporting large coverage and high temporal resolution critical for neuroscientific applicability; 2) it has superior sensitivity, facilitating single-subject analyses and continued increments in spatial resolution; and 3) its limited spatial specificity diminishes precisely in the regime of ultrahigh field and resolution: higher field strength and the higher spatial resolution shifts the weighting of the BOLD response away from large veins toward the spatially specific microvasculature, thus mitigating the primary disadvantage of GE-BOLD (discussed below).

### Sensitivity to large draining veins in GE-BOLD

Several strategies, operating at the analysis-level, have been proposed to mitigate the spatial blurring induced by large draining veins in GE-BOLD. These approaches aim to model and remove venous contributions or to exploit statistical structure in the data to enhance spatial specificity. For example, spatial deconvolution models attempt to estimate and correct for deep-to-superficial draining by ascending veins^88–91^. Phase regression removes correlated fluctuations between magnitude and phase time series that are thought to reflect macrovascular signal^74,92–94^. Multivariate analyses (e.g., support vector machines) exploit the high-dimensional nature of fMRI data to recover fine-scale information up to the nominal voxel size^95^. Differential mapping across analogous but orthogonal conditions can improve the spatial precision of responses by suppressing broad-scale venous effects shared by both conditions (but see Faes et al., 2025)^96^. More recently, Temporal Decomposition through Manifold fitting (TDM) has been introduced to exploit hemodynamic delays for separating macrovascular and microvascular components^97^. While all these methods have enjoyed a certain degree of success, they do not fully mitigate loss in spatial specificity (reviewed nicely elsewhere, see for example^69,74,98–100^).

Here, we took advantage of ultrahigh field, and the related gains in spatial resolution it affords, to reduce spatial blurring related to large draining veins in GE-BOLD, thus improving laminar specificity. First, microvascular weighting increases with field strength due to a supra-linear versus a linear field dependence of the BOLD response amplitude in micro-vs macrovasculature, respectively^62,101^. Second, these supra-linear increases in functional contrast, coupled with the concurrent increases in overall image SNR with magnetic field^62,101^, facilitated higher spatial resolutions, further aided by NORDIC denoising^53,54^ (Figure 1B–C, Supplementary Fig. 2). As predicted previously^53,62^, and supported by our recent^56^ and present empirical findings (Figure 1C, Supplementary Fig. 2, Figure 2A,C-D), additional improvements in microvascular specificity can be expected when voxel dimensions approach the spatial scale of field inhomogeneities generated by large pial veins. In this domain, intravoxel fields become increasingly uniform, thereby diminishing T2*-dephasing around large draining vessels–akin to the mechanism underlying increased quantitative T2*-estimates and reduced signal dropout in T2*-weighted scans at smaller voxel volumes (see Figure 1 and supplementary materials in^56^, respectively). By contrast, microvascular perturbations occur at much smaller scales and remain unperturbed thereby shifting the BOLD weighting toward this spatially specific vascular compartment. Last, higher resolution reduces partial voluming across cortical layers. As a result of these combined factors, we were able to resolve the expected layer IV activation peak in V1 using a single-condition GE-BOLD functional analysis (Figure 2A-B).

Nevertheless, laminar profiles still exhibited superficial bias, underscoring the residual influence of large veins (Figure 2A). At 0.35 mm isotropic voxels, the aforementioned condition of resolution-dependent macrovascular suppression is met only for large pial veins in the CSF (average diameter ∼350 μm)^102^. Further gains in resolution may progressively reduce unwanted sensitivity to smaller pial veins (∼100 μm)^102^. In principle, pushing resolutions beyond ∼100 μm could begin to mitigate unwanted BOLD responses also from the largest ascending intracortical veins (diameters >100 μm)^102^, known to degrade laminar specificity by blurring the BOLD signal across layers (deep → superficial)^58–61^, while maintaining BOLD contrast in capillaries that better reflect neuronal activity.

### Towards true laminar resolution

At conventional submillimeter resolution of approximately 0.5 µL, interpretations are framed in terms of broader layer groups (e.g., superficial, middle, deep) and typically rely on histologically derived estimates of relative layer thickness. This approach carries uncertainty given the sparse number of human laminar histological studies^103–105^, inter-subject variability in laminar proportions^85^, and systematic variation of layer proportions with cortical curvature^76,106^. Thus, a critical step toward functional imaging of individual layers is establishing reliable MR landmarks that link functional profiles to underlying cytoarchitecture, allowing true laminar rather than depth-dependent imaging.

This strategy has been applied in primary motor cortex (M1), where Huber and colleagues combined histology and ex-vivo MRI with ultrahigh-resolution anatomical scans in-vivo to identify laminar markers^23^ that subsequently informed laminar studies of motor disorders^12,13,107^. More recently, building on existing studies imaging the Stria (e.g.^64,66,70,108,109^), Gulban and colleagues showed the Stria was clearly identified as a dip in T2*-weighted laminar profiles using ultrahigh resolution structural scans^65^. In the present study, we extend this approach by going to ∼0.04 µL functional acquisitions, enabling identification of the Stria dip directly from the intensity values of the EPI scans used for functional measurements (e.g. Figure 1A, Figure 2A). This has the advantage of ruling out misalignments of the landmark caused by inaccuracies in co-registration which is particularly challenging at these resolutions (Figure 4, discussed in “*Aligning functional and anatomical scans”*).

Layer IV in V1 was chosen here as a proof of principle for single-layer imaging, given its well-characterized anatomical and functional properties. The concentration of myelinated axons, inducing signal drops in T2*-weighted intensity values, permitted its identification as a dark band within the cortical ribbon. Moreover, interspecies correspondence for this low-level visual area permitted specific predictions arising from the extensive invasive animal literature, thus providing a ground-truth to validate functional maps. It should be noted however that layer IV is amongst the thickest cortical layers, making it comparably easier to resolve. Other layers, such as layer I (∼100 µm), are substantially thinner and will pose greater challenges. Moreover, the Stria only exists within the confinement of V1. To approach true laminar fMRI across a broader set of layers and brain areas, higher resolutions are needed. We view the present results as one such step, with continued resolution-gains remaining a key priority to identify smaller anatomical landmarks outside of V1, such as the internal band of Baillarger^110^.

Nonetheless, the Stria served as a valuable testbed, providing a well-established anatomical marker of layer IV in V1^111,112^. The ability to detect this anatomical microfeature, both functionally and structurally, is critical for true laminar fMRI. It allows precise orientation within the cortical ribbon, through–in this specific case–direct identification of layer IV in V1 (Figure 1A; Figure 2A). Functional peaks can thereby be linked to the underlying layers, facilitating accurate interpretation of laminar profiles (Figure 2F), and preventing reduced effect sizes from averaging misaligned laminar features within or across subjects. It further permits precise layer alignment across ROIs (Figure 3), which is critical at this resolution, and aids fine-tuned alignment (Figure 4; Figure 5). Taken together, these observations support the move towards true laminar fMRI, rather than cortical depth-dependent fMRI, where each depth can only be considered relative to its distance from e.g. the pial surface.

### Challenges associated with ultrahigh field and resolution GE-BOLD

#### Layer positioning

In spite of the benefits discussed thus far, ultra-high spatial resolution comes with additional challenges. One such challenge is that, whilst cortical depth alignment may be relatively forgiving at traditional submillimeter precision (i.e. ∼0.8 mm isotropic)^23,32,113^, ultrahigh resolution protocols leave little margin for error (Figure 3; Supplementary Fig. 4A); even small inconsistencies in layer positioning across the ROI may significantly alter laminar response profiles (<0.5 mm; Supplementary Fig. 4A). However, layer-specific anatomical microstructures in mean-EPI images, such as the Stria, become clearly detectable at these resolutions and can be utilized as a tool to assess the quality of alignment across slices and columns within the ROI (Figure 3B-C; Supplementary Fig. 4B). This reveals inconsistencies in both through-plane and in-plane dimensions and provides a means to refine layering implementations (e.g., local segmentation boundary adjustments). Combined with tools such as 2D histograms and spatial filters for boundary enhancement^114–116^, this strategy supports the high-quality segmentation required to reliably capture layer-specific activation.

The observation that small layer misalignment can significantly impact individual laminar profiles, potentially eliminating highly localized responses, such as the peak of activity in the middle layers of V1 (Figure 3, Supplementary Fig. 4A), indeed represents an additional challenge arising at these resolutions. However, the small tolerance for misalignment at 0.35 mm is concurrently a testament to the spatial precision of the BOLD response at these resolutions.

#### Aligning functional and anatomical scans

An additional challenge that accompanies ultra-high field and resolution is represented by the alignment between functional and anatomical data–a preprocessing step critical for facilitating high-quality segmentation, particularly in regions where the structural CNR of functional images is low.

Whilst complications in co-registering across-modality images with differing FOV, contrast, and distortions are not unique to ultrahigh field/resolution^77,117^, they are greatly amplified in this regime (Figure 4). We found the following factors critical:

1. **Geometric distortions** – EPI acquisitions are nonlinearly warped relative to the minimally distorted ME-GRE reference (Figure 4A). These distortions arise from B0 inhomogeneities that obstruct spatial encoding along the phase-encoding direction. Their severity increases with field strength, and their effect becomes more pronounced with the longer echo spacings associated with higher spatial resolution imaging^78^.
2. **Signal loss** – Regions near air–tissue interfaces and large deoxygenated veins exhibit severe B0 inhomogeneities, leading to dropout^62^. Dropout typically varies across EPI and anatomical scans, because of differing readouts and phase encode directions, which further complicates registration.
3. **Resolution-related image degradation** – At 0.35 mm, EPI data are particularly vulnerable to artifacts such as fold-over due to partial imaging slabs, aliasing artifacts due to parallel imaging, motion-related ghosting, gradient nonlinearity–induced “fuzzy ripples”, and B1 transmit inhomogeneity^77,118–120^.

Previous laminar studies have successfully addressed the milder distortions encountered at lower field strengths and resolutions (≤7T/∼0.8 mm)^121^, where accuracy demands are less stringent. A common approach is distortion correction in preprocessing using estimated B0 field maps obtained from multi-echo or reversed phase-encode acquisitions^77,121^. The latter approach has recently been favored in direct comparisons^122^, and was adopted here as an intermediate step for the co-registration (see *Methods*). While this alleviated global distortions, local distortions remained (Figure 4B). This resulted in misalignments in these regions that could not be compensated by non-linear registration (Figure 4C), which were even exacerbated after this approach in some cases (see *Methods*). In contrast, alignment was precise to the level of the Stria in well-behaved regions (minimal residual distortion, dropout, and artifacts; Figure 4D).

This exemplifies how conventional strategies may fall short at ultrahigh field and resolution, warranting approaches tailored to this precision-demanding regime. Promising alternatives include acquisition of anatomical scans distortion-matched to the functional EPIs^117,121,123,124^, or a brief EPI scan acquired with matched distortions and relevant parameters, but using, e.g., a multi-shot, dual-polarity EPI readout^118^ to serve as a high-quality template for co-registration with the anatomical scan. Looking ahead, continued advances in hardware and acquisition are expected to further mitigate these challenges (see *Limitations and future developments*).

#### Motion correction

Geometric distortions are dynamic over the course of an imaging session, varying with motion, respiration, and B0 drifts^77,81^. The resulting non-linear spatial misplacements are not accounted for with rigid-body motion correction adopted in virtually all existing laminar fMRI studies—perhaps because these have not been a major limitation at ≤7T and conventional laminar resolutions. At ultrahigh field strength and resolution, however, the greater sensitivity to distortions may call for additional non-linear transforms during motion correction.

Indeed, combining rigid and non-linear transforms improved alignment across runs in three of four subjects. No significant difference was found in the remaining subject (Subject 1, Figure 5A), likely due to minimal overall motion related RMS error in this subject (Figure 5C). However, as highlighted by L. Huber (ISMRM 2024, “Ultrahigh spatial resolution imaging in the presence of motion”), non-linear algorithms risk confusing paradigm-driven periodic EPI signal fluctuations (i.e., BOLD activation) with motion, thereby introducing spurious “corrections” that can suppress true BOLD responses—as also observed here (Figure 5B). Accordingly, we restricted non-linear transforms to across-run alignment (for which this is not a concern as they carry no functional information) and relied on rigid alignment within runs. An alternative potential strategy to handle volume-to-volume spatial non-linearities is dynamic distortion correction as proposed by^81^, though this remains untested at ultrahigh resolution.

In our data, rigid alignment resulted in overall consistent volume-to-volume alignment based on the overlap in single-volume EPI profiles (Figure 5C) and motion videos (Supplementary Fig. 6). However, exceptions to this were observed in cases of extreme motion relative to the voxel size (>2 mm), as seen in the two runs with the largest RMS error in Subject 4 (green box in Figure 5C). Motion videos indicated that on top of spatially non-linear misalignments uncaptured by the rigid registration, these runs were additionally corrupted by background fluctuations related to motion and physiological noise (Supplementary Fig. 6B). Such runs are likely better discarded, a conclusion supported by the corresponding functional laminar profile, which deviated markedly from that of the remaining runs (Figure 5D). Consistency checks of single-volume EPI profiles (Figure 5C) may provide a practical means to flag compromised runs independently of functional results.

### Limitations and future developments

EPI acquisitions inherently involve trade-offs between coverage, temporal resolution, and spatial resolution. In this study, spatial resolution was pushed to unprecedented levels, limiting coverage to 40 through-plane slices corresponding to a slab thickness of 14 mm. Temporal resolution was similarly constrained, with a volume acquisition time just under 5 seconds, compatible with task-based fMRI, albeit, in its current form, restricted to block designs.

A common denominator for those limitations is the prolonged readout associated with high resolution acquisitions. Long readouts further underlie the distortions and signal dropouts discussed in *Aligning functional and anatomical scans* and introduces T2*-blurring resulting in an effective resolution below the nominal 0.35 mm along the in-plane phase encode direction. Partial Fourier sampling can compensate for some of these effects to an extent but also adds blurring, particularly when using vendor-default rather than more resolution preserving implementations (Figure 1B). Moving forward, however, hardware advances, including high performance head-gradients and high channel-count receive coil arrays^33,51,55^, promise dramatically accelerated readout speeds. Combined with in-plane segmented (multi-shot) acquisitions^70,125,126^, these ongoing developments have already enabled ∼0.04 μL BOLD and ∼0.06 μL VASO acquisitions at 7T^33^.

Furthermore, no through-plane undersampling was employed here, which could readily be implemented to increase coverage and temporal resolution^125,127^, facilitating temporal resolutions compatible with event-related designs. This is similarly attainable by use of 2D multiband imaging^128^, which does not reduce the amount of sampled data, though imperfect excitation profiles remain a key limitation that must be addressed to achieve sufficiently thin slices for ultrahigh resolution 2D imaging^33^.

Notably, while faster readout (higher bandwidth) and k-space undersampling raise thermal noise levels, this is not currently a bottleneck given the SNR and CNR gains afforded by 10.5 T and NORDIC denoising.

## Conclusion

Understanding brain function in health and disease requires access to the fundamental units governing mesoscopic information flow—namely, the cortical layers. Given the spatial scale at which individual layers operate, this evidently requires higher spatial resolution than the conventional ∼0.5 µL voxel volumes. Crucially, such resolution must be achieved at sufficient fCNR to support single-subject analyses, allowing characterization of individualized percepts and disease phenotypes that may ultimately guide tailored interventions. GE-BOLD, with its high sensitivity and synergy with ultrahigh field, is well positioned to meet these demands.

Here, we demonstrate that GE-BOLD fMRI at 10.5 T and ∼0.04 µL resolution enables in vivo identification of cortical landmarks and functional activation patterns that approach the scale of individual layers in the human brain. By resolving layer IV activation in V1 as identified via the Stria of Gennari, we showcase how ultrahigh field and resolution imaging can narrow the gap between human noninvasive studies and invasive animal work. Leveraging the increased fCNR afforded by ultrahigh field, tailored image reconstruction, and denoising, this was achieved robustly at the single-subject level.

Furthermore, we highlight the methodological challenges that arise in this regime, including distortion, alignment, and layer positioning, and discuss strategies to mitigate them. These issues are expected to become increasingly critical as the field advances toward thinner layers and higher-order cortical regions, beyond the relatively thick layer IV in V1 used here as proof of principle.

Taken together, we believe these findings mark a step toward extending laminar fMRI beyond broad layer groups, aiming to probe the circuitry underlying human brain function at the scale of individual layers.

## Acknowledgements

This work was supported by NIH grants R01 NS136490, NIBIB P41 EB027061, P30 NS076408, S10 RR029672. We thank Alexander Bratch and Matt Waks for scanner support, Alireza Sadeghi Tarakameh for B1 shimming, and Theresa Whitney for recruiting participants.

## Methods

### Participants

We acquired data from four healthy subjects with normal or corrected vision (age range: 28-52 years). Two subjects (Subject 1 and Subject 2) were scanned in two sessions separated by more than a week to showcase replicability. Subject 1 was additionally scanned at 7T for comparison with a “standard” 0.8 mm isotropic acquisition. All subjects provided written informed consent. The study complied with all relevant ethical regulations for work with human participants. The local IRB at the University of Minnesota approved the experiments.

### Imaging protocol

All 10.5 T MRI data were collected with a Siemens Magnetom System (Siemens Healthineers, Erlangen, Germany) with an in-house built 16-channel transmit and 80-channel receive head coil, characterized extensively in terms of performance^51,55^. The 0.8 mm isotropic 7T fMRI data were collected on a Siemens Magnetom System with a single transmit and 32-channel receive NOVA head coil (NOVA Medical, Wilmington, MA, USA).

3D ME-GRE (6 echoes) was used as an anatomical reference to guide segmentation and to identify the Stria. Parameters were: Voxel size: 0.37 mm isotropic; TR=29 ms, TEs: 4.1, 7.9, 12, 16, 20, 23 ms, FA=10deg; 7/8ths PF; in-plane and through-plane GRAPPA: 3×3; Matrix size=576×576×320.

The 3D GRE-EPI sequence used for functional acquisitions at 10.5T had parameters: Voxel size: 0.35 mm isotropic; TR=113ms; TE=24ms; FA=15deg; iPAT=4; 5/8 PF; Matrix size=364×260×40 (40 coronal slices); VAT=4972 ms.

The 7 T functional dataset acquired at 0.8 mm isotropic resolution in one subject using a 2D GRE-EPI readout had parameters: TR=1350 ms; Multiband factor 2, iPAT 3; 6/8ths PF; TE=26.4ms; FA=70deg; Matrix size=198×210×42.

### Experimental design

Visual stimuli consisted of center (target) and surround square checkerboards, counterphase flickering at 6 Hz and subtending approximately 5.6° of visual angle^53^. Stimulation blocks (24 s, target or surround) alternated with 24 s blocks of fixation on the red dot at the center of the screen. Stimuli were displayed on a gray background with an average luminance of 25.4 cd/m² (range 23.5–30.1 cd/m²). At 10.5 T, stimuli were presented on a VPIXX projector (1920 × 1080 resolution at 60 Hz; viewing distance ∼89.5 cm) using a Mac Pro computer. At 7 T, stimuli were presented on a Cambridge Research Systems BOLDscreen 32 LCD monitor positioned at the head (1920 × 1080 resolution at 120 Hz; viewing distance ∼89.5 cm). Stimulus presentation was controlled using Psychophysics Toolbox^129^ (version 3.0.15). Participants viewed the stimuli through a mirror placed in the head coil.

Each run began and ended with a fixation block and comprised three trials per condition (pseudorandomized order). This yielded a run duration of just over 5 minutes (6–8 runs per subject). Participants were instructed to minimize movement and maintain fixation on the central dot throughout the run.

### Tailored offline reconstructions

Achieving an echo time near the T2* of gray matter at high resolution and ultrahigh-field strength necessitates high acceleration factors^130^ and Partial Fourier^131^ (PF) sampling along the in-plane phase-encoding direction (here: PF = 5/8, IPAT = 4). Accelerated imaging increases thermal noise levels which are already substantial at this resolution^132^, and the default vendor implementation of PF can introduce excessive spatial blurring. To overcome this limitation, images were reconstructed offline with a tailored Projection Onto Convex Sets (POCS) implementation of PF^133^ to reduce resolution loss, before being NORDIC-denoised for thermal noise suppression^53,54^ (see Figure 1B and Supplementary Fig. 1 for a comparison with the vendor implementation).

The 3D-EPI data were obtained with monotonic sampling of the slice-phase encoding direction, and preceding each EPI readout a 3-line navigator was acquired without any phase-encoding. Each temporal multi-channel data sample was decorrelated based on the correlation estimate from a noise-only acquisition. All EPI data were processed with a three-step phase correction scheme: 1) even/odd readout correction using linear time correction estimated from a 3-line navigator using opposite polarities; 2) correction for apparent off-resonance motion from B0 (DORK) estimated from the 3-line navigator using same polarity; 3) correction of apparent shot to shot bulk phase variations using a data self-consistency technique. Phase-encoding under sampling was corrected using a two-step GRAPPA implementation, where firstly the missing lines were calculated, and then the acquired data were re-estimated from the neighboring measurements, analogous to a 1-step SPIRIT. POCS partial Fourier reconstruction was applied for each channel and each slice, and the reconstructed images were combined using a SENSE-1 algorithm with ESPIRIT estimated sensitivity profiles. NORDIC denoising was applied to the complex valued time-series as previously described^54^, where the g-factor and noise were estimated using an implementation of the MPPCA algorithm^134^. Magnitude data were used for further processing.

### Preprocessing of fMRI data

#### Preprocessing was kept minimal and performed using AFNI^80^, ANTs^135^, and ITK-SNAP^136^

Functional time series were motion corrected in two steps. First, rigid-body parameters (three translations, three rotations) aligning each volume to the first volume within its run were estimated with AFNI’s *3dvolreg*-function. Second, parameters to align the mean of each run to a reference run (the one closest in time to a reverse phase-encoded scan for distortion correction, see below) were estimated using ANTs’ *antsRegistration*-function, including both linear and non-linear (SyN^135^) transforms. Motion estimation in both steps was restricted to a brain mask. All transforms were then concatenated and applied in a single interpolation step using AFNI’s *3dNwarpApply*-function. This approach (non-linear across-run and linear within-run motion correction) was used for the main analyses. It was compared with a linear across-run registration alternative, implemented by omitting the non-linear transforms during concatenation (Figure 5A). In addition, it was evaluated against non-linear within-run registration, implemented by estimating both linear and non-linear parameters for aligning all volumes in a run to its first volume using *antsRegistration*, followed by reslicing with either the combined linear+non-linear or linear-only transforms (Figure 5B). Note that linear across-run and within-run correction was sufficient for high-accuracy alignment in the 7T 0.8 mm data where motion-induced distortions are less pronounced, particularly for this well-trained subject.

Co-registration between the ME-GRE image (averaged across runs for subjects with two acquisitions and averaged across all echoes) and distorted EPI space proceeded as follows. To bring ME-GRE and mean-EPI images into closer spatial correspondence prior to alignment, a distortion-corrected mean-EPI template was computed from a reverse phase-encoded scan (average of 10 volumes) using AFNI’s *3dQwarp*-function (blip-up/blip-down correction)^79^. The parameters to align the ME-GRE image to this template were then estimated using *antsRegistration* (rigid and non-linear transforms). To minimize interpolation-induced blurring of functional images through the distortion-correction step, the inverse of the distortion deformation field was computed, concatenated with the “ME-GRE → distortion-corrected EPI” transforms, and applied to align the ME-GRE to distorted EPI-space in which all further analyses were performed. Note that in two subjects, the ME-GRE was instead aligned directly to the non-distortion-corrected mean-EPI which yielded more accurate alignment locally around the ROI in those cases.

Activation maps were computed from motion-corrected time series using a standard general linear model (GLM)-framework^137^ (AFNI’s *3dDeconvolve*-function). Time series were scaled independently per run to have a mean of 100, yielding beta-estimates expressed in PSC. The design matrix consisted of a boxcar paradigm-regressor (trial onsets and offsets) convolved with a double-gamma function and polynomial regressors (up to the third order) for regressing out low-frequency fluctuations. To compute across-trial standard error of laminar profiles, an additional GLM was run to obtain single-trial beta estimates using one paradigm regressor per trial per run.

### ROI definition and segmentation

The ROIs for laminar analyses were defined in the retinotopic representation of the target condition within V1. First, V1 was localized based on the Benson probabilistic atlas^138^. The target representation was then determined from robust target-preference based on target>surround contrast maps. Within this region, the ROI was further restricted to a cortical patch (spanning multiple slices) that exhibited sufficient anatomical contrast for segmentation and a clear Stria.

Voxel-wise cortical depth estimates ranging from 0 to 1 (WM to CSF) were obtained with LAYNII (*LN2_LAYERS*-function), using manually drawn WM and CSF boundaries. These were drawn around the ROI on an upsampled 0.2 mm isotropic grid, guided by the mean-EPI and ME-GRE images. Laminar profiles were generated by binning the cortical ribbon into 13 depth ranges followed by averaging of voxel values (e.g., image intensity or functional PSC) within each bin. The ROI was further divided into “columns” (*LN2_COLUMNS*-function) (Figure 3A) used to assess layer positioning (see below).

To ensure consistent layer positioning, ROI segmentation was refined iteratively: Starting from boundaries drawn from the anatomical contrast, we plotted the resulting laminar mean-EPI profiles of each slice (through-plane) and column to also capture in-plane consistency (Figure 3A). This was repeated for different layering parameters, i.e. equidistance versus equivoluming and varying smoothing iterations (more iterations shift equivolume estimates toward equidistance). The optimal implementation was determined from overlap of depth-dependent features in mean-EPI laminar profiles (including the Stria dip) across slices and columns (Figure 3B-C). Residual inconsistencies were corrected by adjusting the segmented boundaries on a slice-by-slice or column-by-column basis, informed by these plots.

The slice-to-slice misalignment of the exemplar inconsistent layer positioning shown in Figure 3B was quantified as the difference in relative depth between the local minima in each slice-wise mean-EPI intensity profile and that of the across-slice mean profile. These depth differences were then converted to millimeters using the average cortical thickness of the ROI estimated by *LN2_LAYERS*. To further illustrate how misalignment of depth estimates may impact functional laminar profiles, we implemented systematic shifts of depth maps relative to the activation maps, mimicking inconsistent layer-positioning across the ROI. Specifically, data of each slice was upsampled from 0.2 mm iso to 0.05 mm iso using nearest neighbor interpolation, and the planes of voxel-wise depth estimates were then shifted along the in-plane dimension having the largest impact on slice-wise mean-EPI profiles. This was repeated for a range of shift sizes (±0.1-0.3mm, corresponding to total between-slice misalignments of 0.2-0.6 mm) and for all possible shift patterns across slices, with the constraint that positive and negative shifts were balanced. Functional laminar profiles were then computed from the unshifted and shifted depth maps and averaged across slices (Supplementary Fig. 4).

### Statistical analyses

To assess statistical significance of the functional activation peak at the depth of the Stria in laminar profiles (Figure 2), we applied the following procedure independently per subject. For every trial, we fit a model consisting of a second-order polynomial (capturing the characteristic ramping profile of GE-BOLD), plus a Gaussian term (capturing the localized activation peak). Fitting was performed using MATLAB’s *lsqcurvefit* function (MathWorks Inc.). The model comprised three polynomial parameters (intercept p_1_, slope p_2_, quadratic term p_3_) and three Gaussian parameters (amplitude p_4_, mean p_5_ (depth-position), standard deviation p_6_ (width)). To prevent physiologically implausible solutions arising from overfitting—such as a monotonic decreasing polynomial compensated by an inflated Gaussian peak—we employed a leave-one-out fitting strategy. Specifically, for each trial, the polynomial component was first fitted to the mean profile of all remaining trials (polynomial-only model). The resulting slope (*p₂*) and quadratic term (*p₃*) were then fixed in the subsequent full model fit (polynomial + Gaussian; Equation 1) for the held-out trial. This procedure reduced model flexibility and was physiologically motivated by the assumption that the overall ramping shape of the GE-BOLD profile is largely constant across trials.

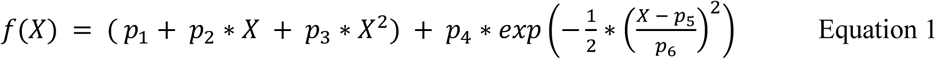

The intercept, slope and quadratic term were unconstrained. The Gaussian mean was constrained to the depth-bin range (1-13), and its standard deviation was constrained to the range corresponding to FWHM=2-6 bins to avoid modelling of physiologically implausible peak widths. The Gaussian amplitude, which constituted the parameter of interest, was unconstrained; the activation peak was considered statistically significant when this parameter was greater than zero, evaluated using a one-sample, one-sided t-test across trials. This directional choice was motivated by extensive prior literature establishing a strong *a priori* expectation of a positive activation peak at middle cortical depths^36–38,63,67,68^; deviations in the opposite direction would therefore constitute evidence against, rather than support for, the ability of the present setup to resolve feedforward layer IV activity.

The minimum shift size of depth maps relative to functional activation maps that resulted in a significant impact on laminar profiles (Supplementary Fig. 4A) was determined as follows. For each trial, the same modelling approach described above was first applied to the unshifted data. Coefficients of determination (R^2^) were then computed between this fit and both the unshifted (for all trials) and shifted (for all trials and shift patterns) profiles. Statistical significance of a given shift size was determined using a bootstrap-derived 95% confidence intervals across trials of R^2^_unshifted_ minus the median R^2^_shifted_ across shift patterns (the median and bootstrapping were used due to the bounded and non-normal distribution of R^2^). To control the family-wise error rate within each subject, tests were performed hierarchically, starting with the largest shift (±0.3mm), and subsequent step-sizes were only tested if the preceding larger shift reached significance (fixed-sequence multiple comparisons correction^139^).

To evaluate alignment accuracy, we examined single-run EPI profiles for across-run assessment (Figure 5A) and single-volume EPI profiles for within-run assessment (Figure 5C). Profiles were z-score normalized across depths to control for low-frequency intensity fluctuations (relevant at both the run and volume levels) and for BOLD-related intensity changes (relevant for single-volume profiles). Alignment accuracy was quantified by subtracting each profile (single-run or single-volume) from the corresponding mean profile (average across runs or volumes), yielding one error value per profile per depth. The RMS error was then computed across profiles for each depth and subsequently averaged. Differences in RMS error between linear and non-linear across-run alignment were tested independently per subject using paired two-sided t-tests. Note that for Subject 1, Subject 2 and Subject 4, five additional EPI runs acquired in the same session with identical acquisition parameters, but using a different paradigm, were included in this analysis to increase sample size (not available in Subject 2).

The significance level was set at α = 0.05.

