## Supplementary Figures for "Bridging the gap with invasive imaging: promises and challenges of a new generation of ultrahigh resolution fMRI"

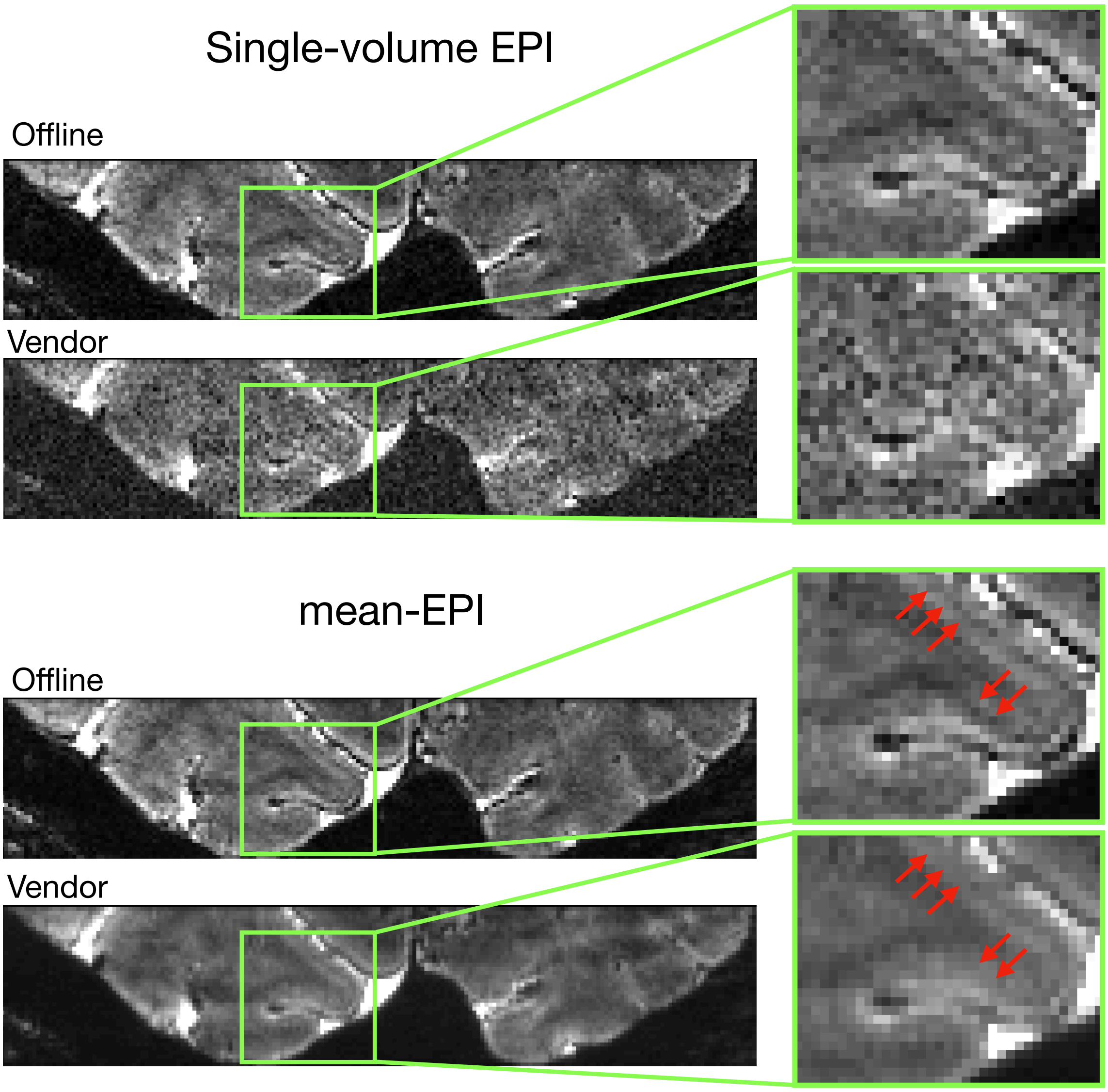


**Supplementary figure 1.** Comparison of single-volume and mean-EPI images reconstructed with either the default vendor implementation or the offline approach. Thermal noise is strongly reduced in the offline reconstruction which included NORDIC denoising, making the Stria visible even in single-volume EPI images. Based on the mean-EPI images, blurring was reduced in the offline reconstruction, yielding a more clearly defined Stria as highlighted by red arrows.

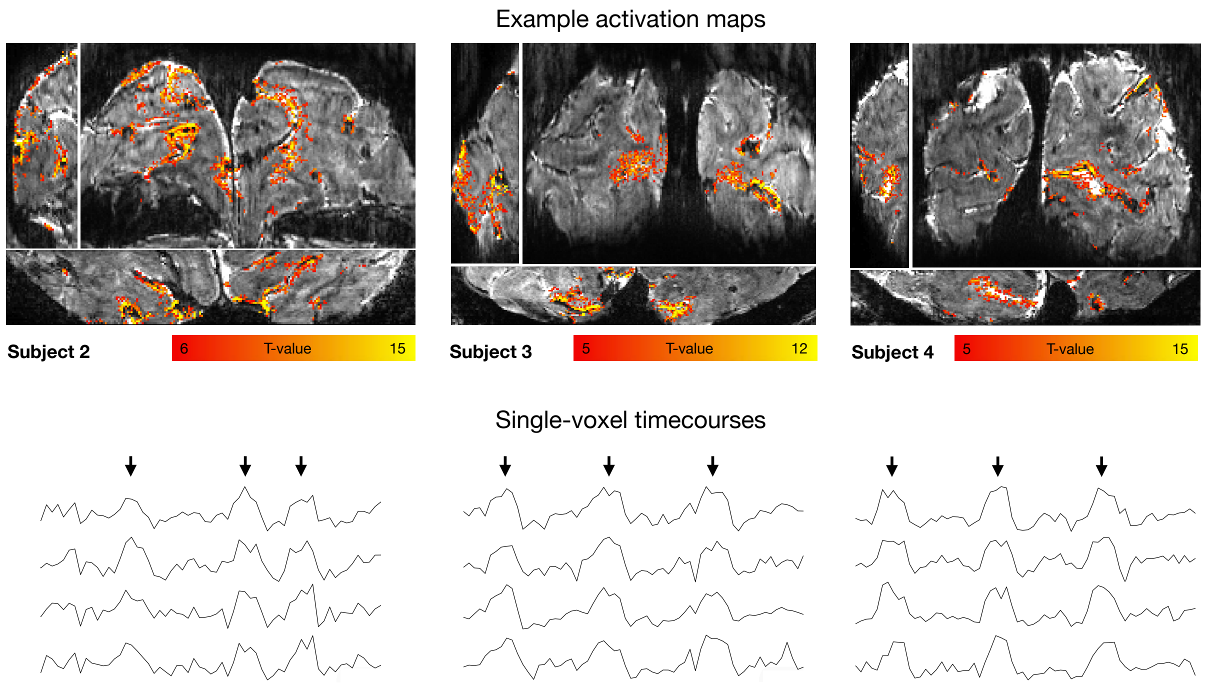


**Supplementary figure 2.** Example activation maps (t-values, target > fixation; upper panel), and single-voxel, single-run time courses (lower panel; black arrows point to stimulation blocks) as shown for Subject 1 in Figure 1C.

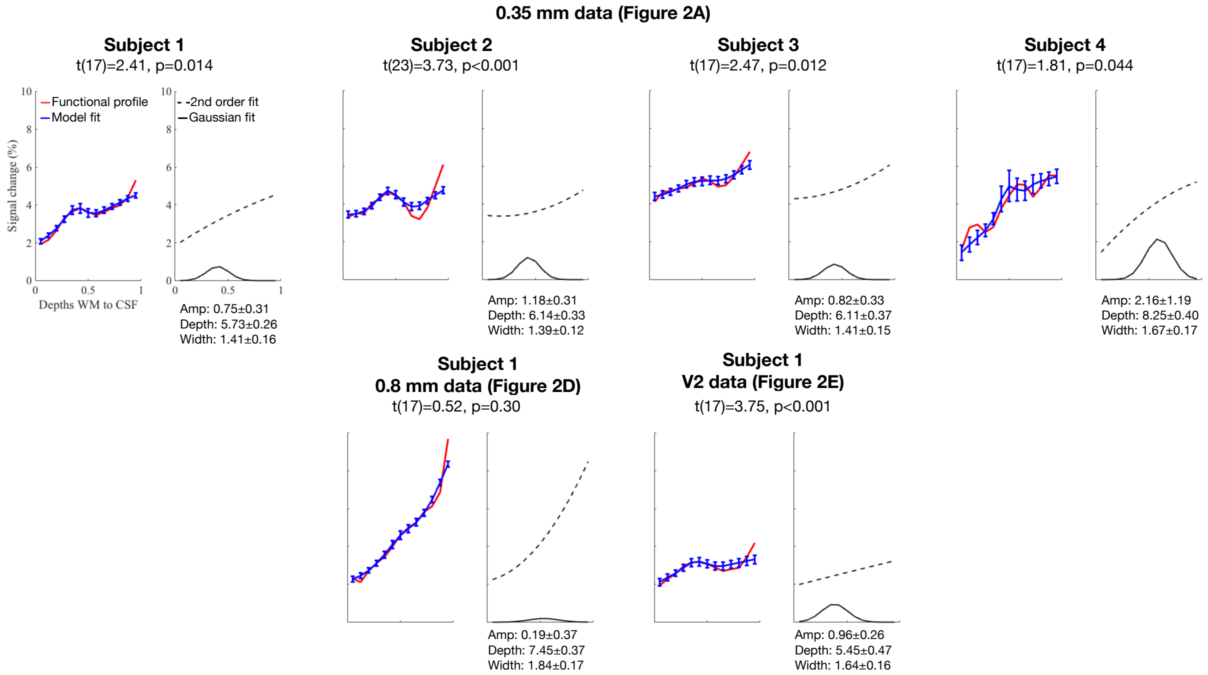


**Supplementary figure 3.** To test whether the activation peaks observed at the depth of cortical layer IV (Figure 2) were statistically significant at the single-subject level, we fit a model comprising a second-order polynomial term (capturing the ramping profile of GE-BOLD) plus a Gaussian term (capturing the peak) to the functional profile of each trial (see *Methods*). Left panels show empirical laminar profiles (red; Figure 2) overlaid with the mean±SE model fits across trials (blue). Right panels show the isolated contributions of the polynomial (dashed) and Gaussian (solid) components, computed from the trial-averaged parameter estimates (mean±SE of the Gaussian amplitude (amp), mean (depth) and standard deviation (width) are reported below each panel). The amplitude was significantly larger than 0 in all 0.35 mm V1 datasets (upper row), as well as in the 0.35 mm V2 dataset (bottom right), and not in the 0.8 mm V1 dataset (bottom left). Importantly, the polynomial components were monotonically increasing, and the estimated depth-position of the Gaussian deviated on average by less than half a depth bin from the peak observed at the Stria in empirical profiles (mean±SE across 0.35 mm datasets: 0.42±0.14 bins), suggesting that the polynomial and Gaussian terms modelled the ramp and peak, respectively, as intended.

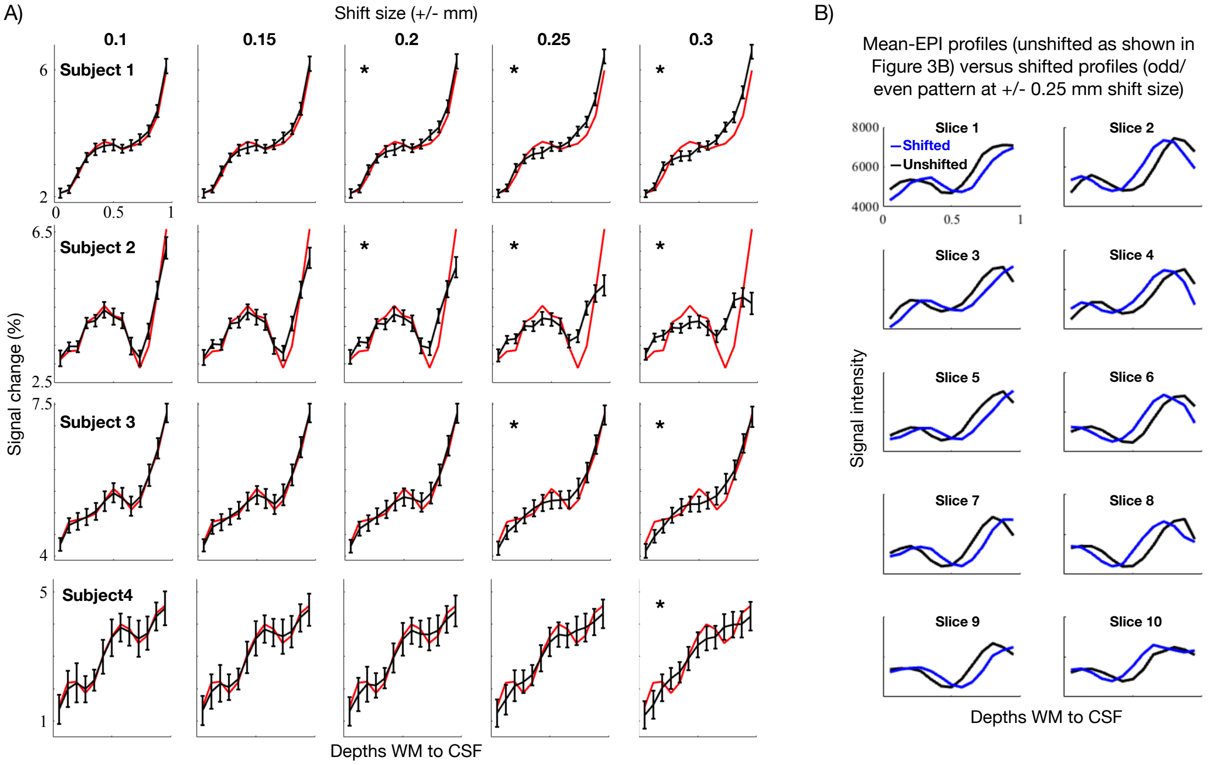


**Supplementary figure 4.** Impact of inconsistent layer positioning on laminar profiles. To mimic inconsistent depth estimates across the ROI, we applied systematic in-plane shifts of the voxel-wise depth estimates across slices (see *Methods*). **A)** Functional laminar profiles (black) are depicted following shifts of $\pm$0.1-0.3 mm (averaged across slices and all possible balanced shift-combinations), overlaid on the unshifted profiles (red) for each 0.35 mm V1 dataset (note that the trials of the first 2 runs were discarded in Subject 2 to keep N=18 for all subjects). Error bars reflect standard error of the mean across trials. Asterisks denote shift sizes at which the shifted profile was significantly different from the unshifted (see *Methods*). The Stria-peak appears gradually less discernible at larger shifts, with the smallest significant shift-size being ±0.24 mm (i.e. a total misplacement of 0.48 mm between slices) on average across subjects. These results support the notion that, at these resolutions, even small inconsistencies in layer-positioning can substantially impact laminar profiles. Importantly, we do not interpret this shift-size as a hard threshold for misalignment effects, as the impact depends on multiple factors such as orientation of the misalignment with respect to the layers, proximity to large veins, cortical thickness, and the structure of misplacements (e.g. random jittering versus systematic shifts). Rather, these results serve to illustrate the necessity of consistent layer-positioning at these resolutions as subtle laminar responses otherwise risk attenuation. As illustrated by the empirical example in Figure 3B–C, mean-EPI profiles can serve as a tool to facilitate such consistency. In panel **B)** we extend this observation in a more controlled manner, by applying systematic in-plane shifts to the same dataset used in Figure 3B and visualizing their effect on slice-wise mean-EPI profiles. Specifically, the unshifted profiles from Figure 3B (consistent, after correction) are plotted together with profiles following ±0.25 mm shifts, corresponding to the across-subject average shift size yielding a significant difference in panel *A)*. Notably, the imposed shift pattern (here positive shifts for odd slices and negative shifts for even slices) is reflected in the shifted profiles, which are displaced toward CSF in odd slices and toward WM in even slices. Observed patterns of misalignment across slice-wise mean-EPI profiles may thereby provide clues for identifying and correcting misalignment, facilitating consistent layer positioning across the ROI.

Corrected forward-EPI versus corrected reverse-EPI

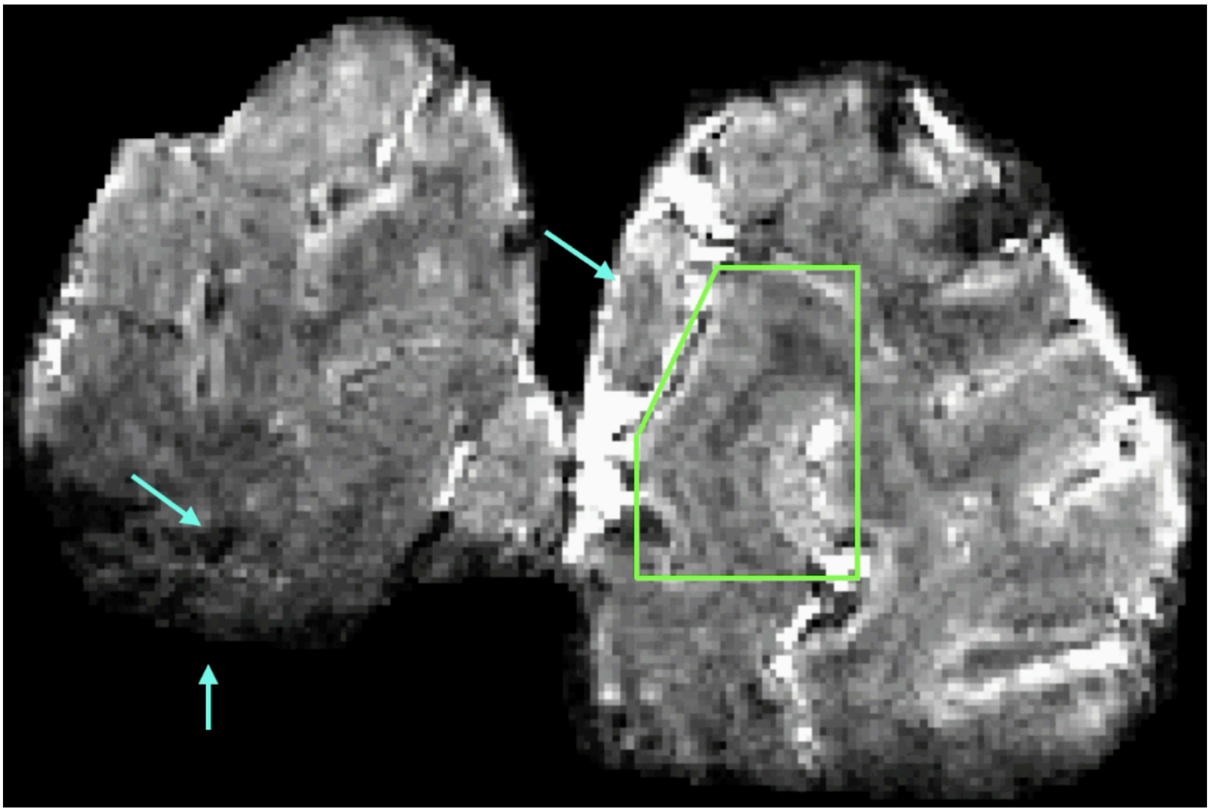

**Supplementary figure 5.** Video alternating between an example coronal slice of the corrected forward-EPI and reverse-EPI images (same as in Figure 4B). The green box highlights a region where residual distortions after distortion correction were small relative to example regions highlighted by the blue errors. In such regions, alignment with the minimally distorted ME-GRE image was generally of high quality, as shown in Figure 4D.

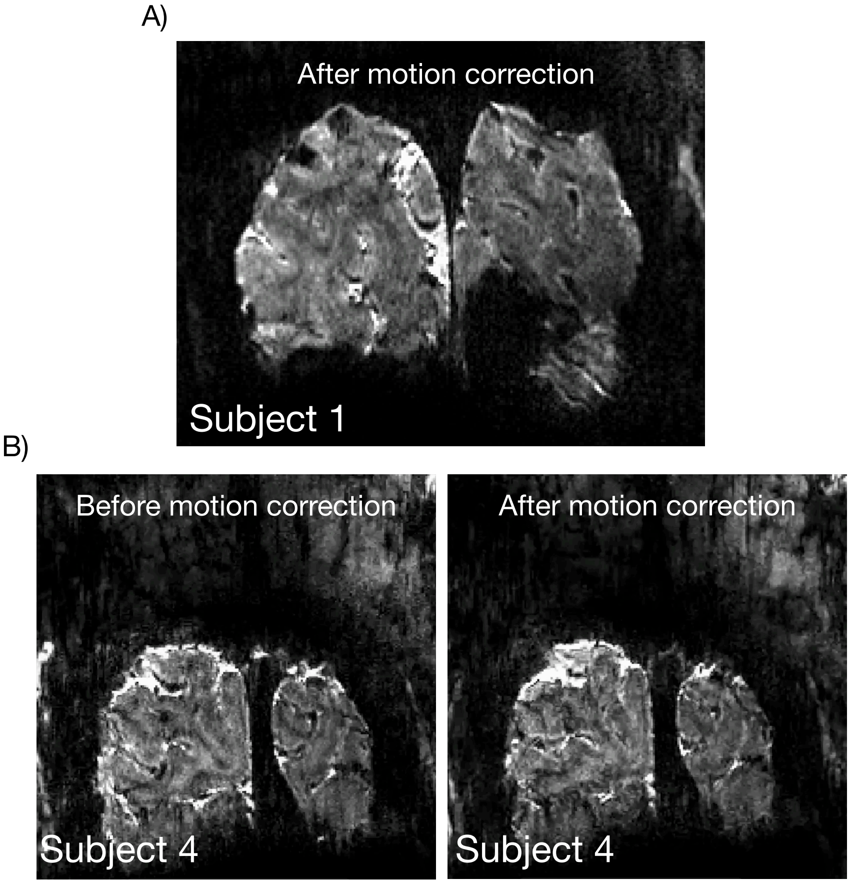


**Supplementary figure 6. A)** Example motion video with relatively little motion (Subject 1 – the run with the highest RMS in Figure 5C), representative of the well-trained subjects recruited in this study. It is played from the beginning to the end of the run to show overall consistent volume-to-volume alignment using rigid-body motion correction (i.e. no non-linear transformations). Video can be downloaded from: 10.6084/m9.figshare.31954419 **B)** Example motion videos of a run with extreme motion (Subject 4 – the second of the two runs highlighted by a green box in Figure 5C). This is shown before and after motion correction (rigid body only, no non-linear transformations). The videos alternate between forward (start → end) and reverse (end → start) playback to highlight: (1) the degree of motion before correction (right); (2) that even for this extreme example, after motion correction, the brain is rigidly well-aligned across volumes but corrupted by background fluctuations moving relative to the now fixated brain. Videos can be downloaded from: 10.6084/m9.figshare.31954419
